# Bacterial dispersal and host specificity shape microbiome strain sharing among great ape species in neighboring zoo exhibits

**DOI:** 10.64898/2026.10.06.757053

**Authors:** Pauline Krauß, Antonia Breier, Anna-Lena Hohlfeld, Hanna Keller, Virginia Kunkel, Max Meier, Sina Plettemeier, Willi Thomas, Reena Debray

## Abstract

Animal microbiomes often exhibit host-species-specific structure, which could suggest a shared evolutionary history with their hosts. However, in wild populations this possibility has been difficult to disentangle from geographic or ecological barriers to microbial dispersal, particularly in taxa like non-human primates in which phylogeny and geography are closely related. To investigate microbiome transmission and adaptation in a common environment, the gut microbiomes of zoo-housed great apes were profiled longitudinally for four months. Even in a shared environment, phylogenetic structure emerged in both species- and strain-level microbiome composition. Longitudinal tracking of new strain acquisitions revealed that transmission across exhibits was possible, even between distantly related host species, suggesting that phylogenetic structure emerged in the longer term as different strains survived in different hosts. Comparison to metagenomes from humans and wild great apes revealed little evidence that human keepers, researchers, or zoo guests drive transmission between exhibits. Instead, across-exhibit transmission disproportionately involved spore-forming bacterial genera, possibly suggesting airborne spore transmission as a means of bacterial dispersal. By investigating microbiomes of different host species in a common geographic environment without direct inter-host interaction, this study identifies spatial constraints on transmission, bacterial dispersal ability, and differential survival of bacterial strains in different hosts as factors shaping the gut microbiomes of great apes.

## INTRODUCTION

As animals evolve and diversify, so do the communities of microbial symbionts that live in, on, and around them. Over time, differences in the physiology, immune activity, and/or diets of different animal species could select for microbes to adapt to particular host species over others, a phenomenon known as host specificity (Mallott and Amato 2021). In wild populations, microbiome adaptation to hosts is often diagnosed based on patterns such as phylosymbiosis or co-diversification. Phylosymbiosis occurs when microbiome community composition reflects the evolutionary history of hosts (Lim and Bordenstein 2020) while co-diversification occurs when closely related microbial lineages associate with closely related host species (Moeller et al. 2023). Some of the clearest examples of phylosymbiosis and co-diversification have come from gut microbiome research in the great apes (Brooks et al. 2016; Sanders et al. 2023; Narat et al. 2020; Nishida and Ochman 2021; Rühlemann et al. 2024). For instance, a study comparing bacterial genomes from humans and 22 additional non-human primate species found significant evidence of host-bacterial co-diversification in 10 bacterial phyla and 206 bacterial lineages (12% of the total lineages tested) (Sanders et al. 2023). Studying microbiome adaptation in great apes may help to manage their welfare in the wild and captivity, and as the closest living relatives of humans, great apes can also help us understand the role that the microbiome has played in our own evolution. For example, investigating which microbial species and genes in the oral microbiome tracked the great ape phylogeny, and which ones diverged from it, revealed rapid adaptation of *Streptococcus* bacteria along the human line to bind salivary amylase after their human hosts evolved to eat and cook starch-rich foods (Fellows Yates et al. 2021).

However, other processes besides adaptation of microbiomes to hosts can produce patterns similar to phylosymbiosis and co-diversification, including environmental correlates of host phylogeny (Mazel et al. 2018) or geographic constraints on bacterial transmission (Good 2026). In some taxa, such as wild mice, it has been possible to sample many wild populations so that phylogeny and geography are not confounded (Dillard et al. 2026). But due to the phylogeographic history of the great ape radiation and the small number of species still surviving today, all members of the Ponginae subfamily live in southeast Asia, while all members of the Homininae family live in Africa, making phylogeny and geography inextricably linked in the wild. Other studies inoculate germ-free hosts with bacteria in the lab to test whether native bacteria are locally adapted (Kwong and Moran 2015; Sprockett et al. 2023), but such methods are not logistically or ethically feasible in great apes.

Zoo-based primate research offers the possibility to study great apes housed in species-specific social groups in a common geographic environment. Zoos, and research on known individuals from habituated social groups in general, also permit repeated sampling from the same individual over time. Longitudinal sampling is critical for distinguishing transmission – which hosts are temporarily exposed to which microbial strains – from adaptation, which determines which strains actually establish and survive in the long term (Debray et al. 2025).

The Zoo Leipzig is home to all four taxa of great apes: chimpanzees, bonobos, gorillas, and orangutans. Chimpanzees and bonobos are the most closely related to each other (diverged approx. 2 million years ago) and to humans (diverged 6-8 million years ago). Chimpanzees are found in west Africa and the Ivory Coast, while bonobos live in central Africa. Gorillas live in central Africa as well, and split from the chimpanzee/bonobo/human lineage 8-10 million years ago. Orangutans are the most phylogenetically and geographically distinct; they live in southeast Asia and split from the other lineages approximately 15 million years ago. The zoo setting thus brings great ape species that do not naturally encounter each other into close proximity. The Zoo Leipzig is especially suited for disentangling bacterial adaptation from other constraints on transmission because there are two separately housed chimpanzee exhibits. Elevated microbiome similarity between chimpanzees living in different exhibits would indicate host-species-specific selection on the microbiome. In this study we combined longitudinal fecal sampling from all five great ape exhibits (one exhibit each of gorillas, bonobos, and orangutans and two chimpanzee exhibits) with shotgun metagenomic sequencing and strain-resolved analysis to characterize microbiome species- and strain-level sharing within and between exhibits.

## RESULTS

### Microbiome composition within and across exhibits

Over a four-month period, keepers at the Zoo Leipzig collected fecal samples opportunistically from known individuals in the five adjoining great ape exhibits (Figure **1a**, **1b**). The samples encompass western lowland gorillas (*Gorilla gorilla gorilla, n* = 14 samples from 6 individuals), bonobos (*Pan paniscus, n* = 15 samples from 7 individuals), western chimpanzees and hybrids of western/eastern chimpanzees (*Pan troglodytes, n* = 17 samples from 11 individuals) and Sumatran orangutans (*Pongo abelii, n* = 12 samples from 6 individuals). We generated shotgun metagenomes and profiled species-level composition using the *kraken2* bacterial database supplemented with 16,141 additional bacterial and archaeal genomes derived from the gut microbiomes of humans and great apes (Sanders et al. 2023; Rühlemann et al. 2024). The gut microbiomes of the apes were dominated by fiber-fermenting taxa such as Lachnospiraceae, Prevotellaceae, and Oscillospiraceae (Figure **1c**). Consistent with previous reports (Narat et al. 2020; Rühlemann et al. 2024), microbiome composition varied across host species, and microbiome compositional differences across species were strongly correlated with the host species phylogeny. Orangutans, the most deeply diverged species, had the most distinct microbiomes from all other animals, followed by gorillas. One gorilla, Kumili, did not cluster with any of her groupmates for reasons that are not known; she was not an outlier in terms of age or sex, and she was genetically related to several of the other gorillas we sampled. Chimpanzees and bonobos, the most closely related species, had the most similar microbiomes (Figure **1d**). The chimpanzee samples formed two subgroups that did not cluster by exhibit. Both of the male chimpanzees in our study (one from group A and one from group B) are found in the upper cluster together with several females, while the lower chimpanzee cluster consisted of only females (Figure **1d**).

**Figure 1:**
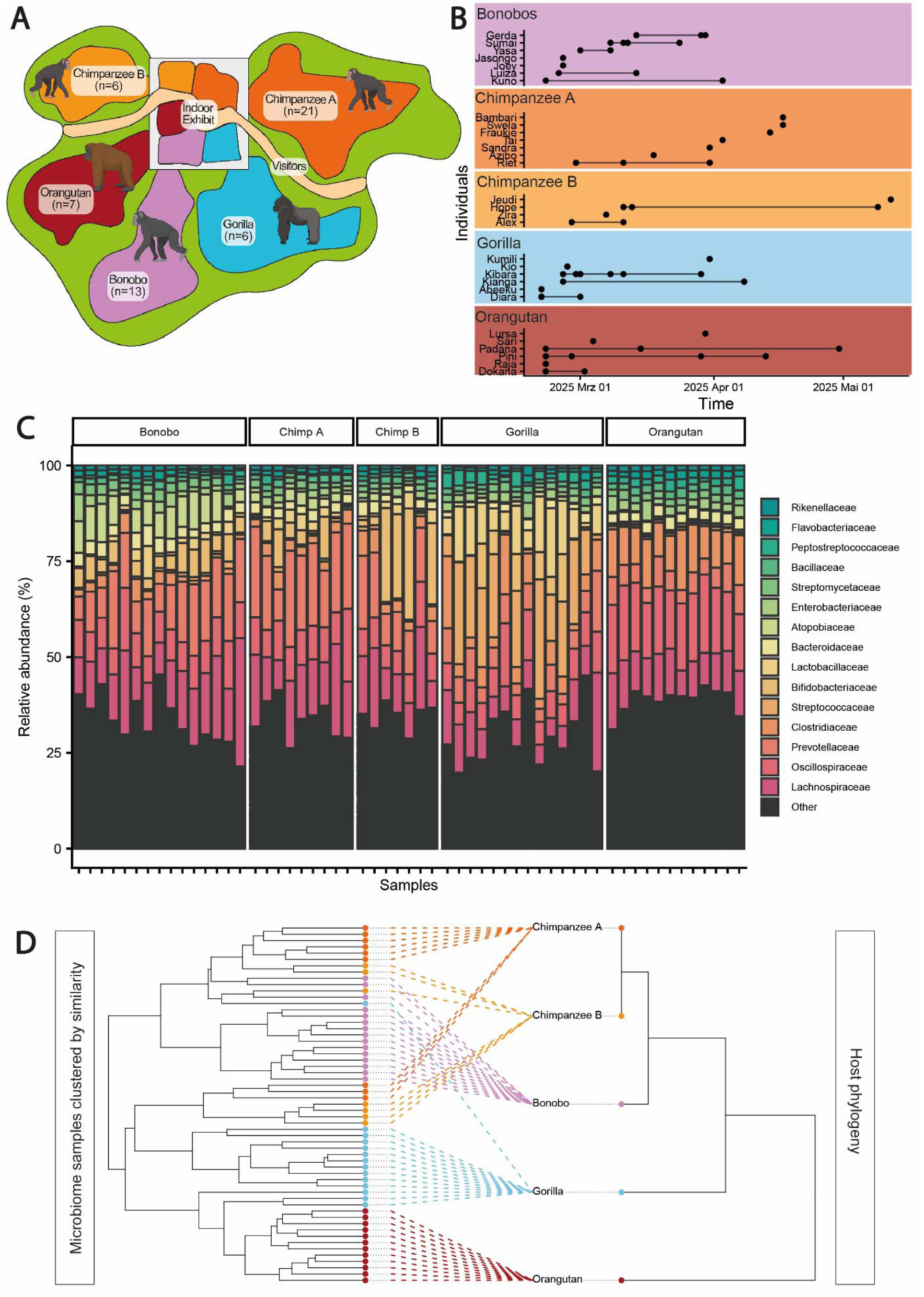
Microbiome species sharing within and across exhibits. **a)** Zoo Leipzig houses four great ape species in five isolated exhibits, each consisting of an indoor and outdoor area. Number of animals in each exhibit is indicated (note that it was not possible to sample all animals). **b)** Sampling overview of opportunistically sampled individuals. Each dot represents one fecal sample; connecting lines link samples from the same individual. **c)** Gut microbiome composition of each collected fecal sample, colored by relative abundance of the 15 most frequent bacterial families. Samples are grouped by exhibit. **d)** Dendrogram of individual fecal samples clustered by microbiome similarity (left), and host species phylogeny (right), both colored by exhibit.

### Strain sharing within and across exhibits

Elevated species-level microbiome similarity among closely related ape species could either reflect convergent selection for similar types of microbes, or shared gut microbial strains carried by multiple animals in different exhibits. To distinguish between these possibilities, we identified instances in which different individuals carried near-identical populations of the same microbial species using *inStrain* (Olm et al. 2021) which measures strain sharing using genome-wide sequence similarity, and *SynTracker* (Enav et al. 2025), which measures gene synteny. In total 1215 strain sharing events between two individuals were observed by combining both pipelines. Strain sharing percentage was calculated by dividing the number of shared microbial strains by the total number of shared microbial species (Figure **2a**).

**Figure 2:**
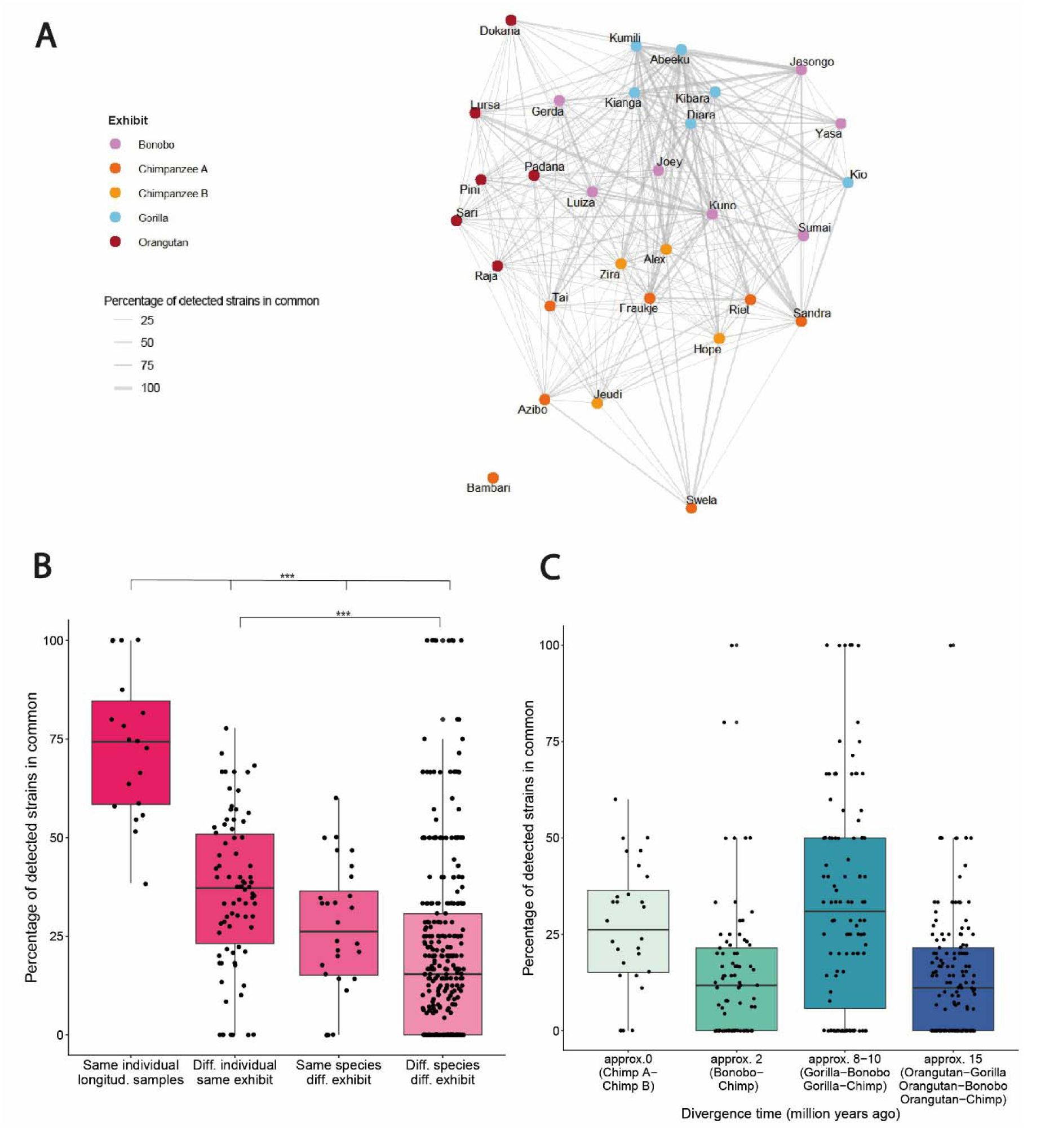
Strain sharing within and across exhibits. **a)** Strain sharing network of sampled individuals. Each dot represents one individual, colored by exhibit. Longitudinal samples were combined for each individual. Lines connect individuals with at least one strain in common. Heavier lines indicate higher percentage of shared strains, where the denominator represents the total number of shared species between the two individuals. **b)** Strain sharing between longitudinal samples, members of the same exhibit, chimpanzees housed in different exhibits, and great apes of different species in different exhibits. **c)** Strain sharing between across-exhibit dyads only. Here, a divergence time of zero years refers to chimpanzee groups in different exhibits, not members of the same exhibit. Overall, divergence times of hosts weakly correlate with strain sharing percentages. (ANOVA, F=3.1449, p= 0.07702). Note that in **b)** and **c)**, a strain sharing percentage of 100% means that for *the subset of species they had in common* their corresponding strains were all near-identical, not necessarily that they had identical microbiomes or identical sets of species.

Strain sharing was highest between longitudinal samples, with individuals typically retaining 50-75% of strains between samples taken a few weeks or months apart. Strain sharing was next highest between individuals of the same species living in the same exhibit, and then between individuals of the same species living in different exhibits (i.e., Chimpanzee A Group members compared to Chimpanzee B Group members). Individuals of different species from different exhibits had the lowest levels of strain sharing (Tukey’s Honest Significant Difference Test, all same-individual comparisons to other categories p<0.001; different individuals in same exhibit vs. different species in different exhibits p<0.001; Chimpanzee A-Chimpanzee B comparisons not significantly different from either different individuals in the same exhibit or different species in different exhibits) (Figure **2b**). Strain sharing rates across different exhibits were weakly correlated with the phylogenetic relatedness of the host species (ANOVA, F=3.1449, p= 0.077) (Figure **2c**). One member of the Chimpanzee A Group, Bambari, appeared not to share any strains with her group-mates. The fact that the Chimpanzee A Group was the least extensively sampled of the five exhibits, and that chimpanzees form subgroups within larger groups (Duncan et al. 2022), suggests that Bambari’s closest social or spatial connections may not be represented in our dataset.

### Tracking strain acquisition events

Total levels of strain sharing are presumably the combined result of which strains animals acquire and which strains they both retain over time, making it difficult to infer transmission patterns from strain sharing alone (Debray et al. 2025). To disentangle these two processes, we used close-in-time longitudinal samples to identify new strain acquisitions in the great ape community that occurred during the four-month study. Similar to previous work tracking strains through wild baboon groups (Debray et al. 2026), we identified new strain acquisitions when our samples met three criteria (Figure **3a**). First, the individual in question had to be sampled at least twice, first without the strain and then with the strain. Second, the new strain must have been previously present in at least one other individual, considered the possible donor. Finally, the recipient sample supporting the initial absence of the strain had to occur either after the possible donor was sampled, or less than seven days before the donor was sampled, to support inference about the directionality of transmission from donor to recipient. If the only samples supporting the initial absence of the strain in the recipient were from much earlier than the samples supporting the initial presence of the strain in the donor, then it would be more plausible that the recipient had actually acquired the strain before the donor (i.e. from some other source). Based on these criteria, we identified 176 transmission events from 68 unique donor-recipient dyads.

**Figure 3:**
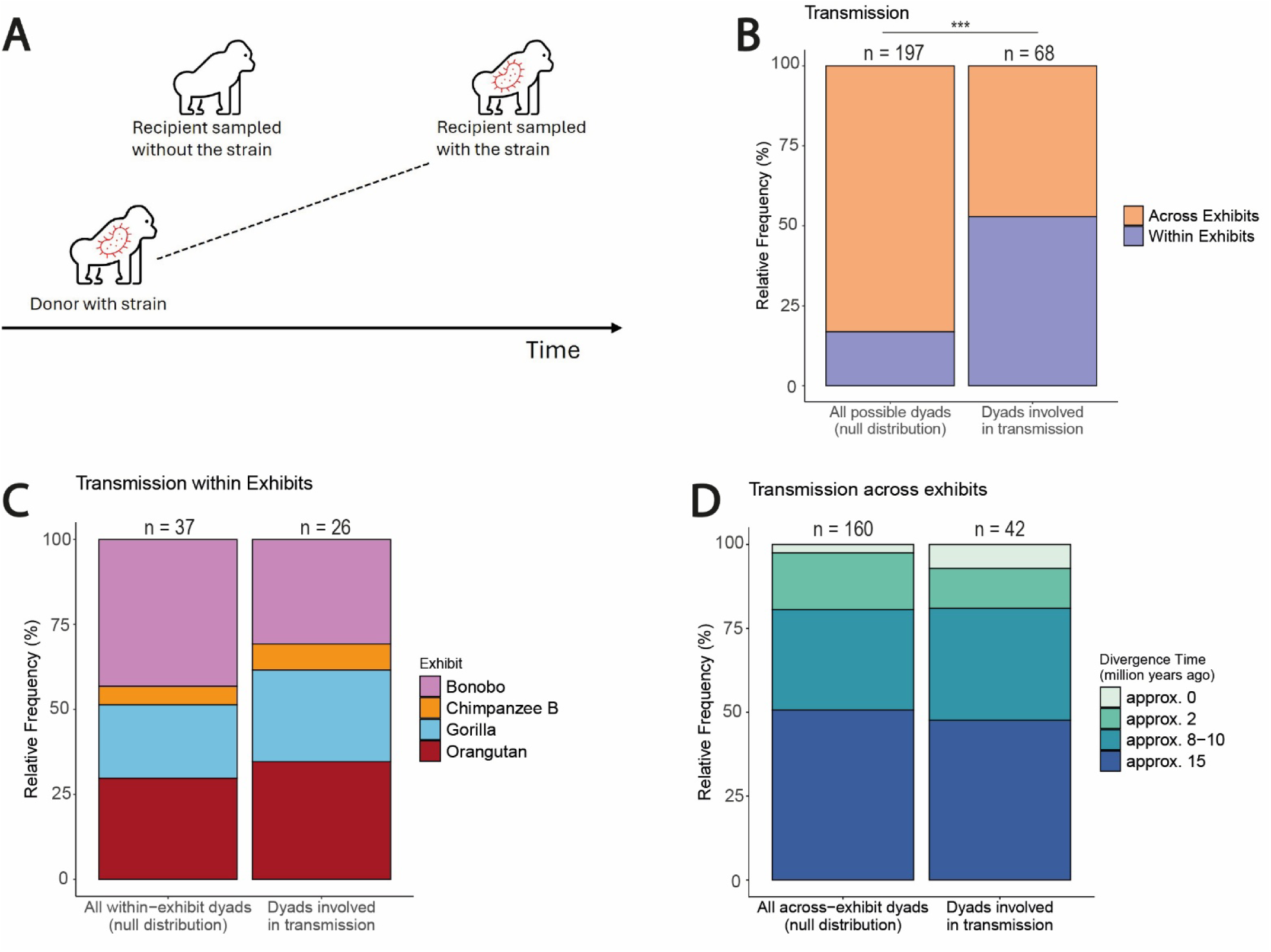
New strain acquisitions during the four-month study period. **a)** We used the following criteria to identify transmission events: one individual (the possible donor) had to be sampled with the strain in question. Another individual (the recipient) had to be sampled twice, first without the strain and later with the strain, such that the strain acquisition was new. The early sample had to be taken either after the donor sample or up to 7 days before. **b)** All animal dyads detected in transmission events compared to a null distribution. Dyads involved in transmission events were more likely to live in the same exhibit (binomial regression, coefficient= 1.8932, p<0.001). **c)** All animal dyads detected in within-exhibit transmission events compared to a null distribution. There were no significant differences in strain transmission within groups (binomial regression; p=0.441). **d)** All animal dyads detected in across-exhibit transmission events. A divergence time of zero years refers to chimpanzee groups in different exhibits. Phylogenetic relatedness of hosts had no significant effect on transmission events across exhibits (binomial regression; coefficient=x, p=0.14984).

Because samples were collected opportunistically, there was variation across individuals and exhibits in the number and timing of available samples. We constructed a null distribution of all animal pairs for which we had the appropriate number and timing of samples to detect transmission according to our criteria if it had ever occurred. For instance, we should be able to detect strains transmitted between the orangutans Padana and Pini, because each individual had a sample that was collected before at least two subsequent samples from the other (Figure **1b**). Conversely, it was not possible to detect any within-group transmission among members of the Chimpanzee A Group, because only one member was sampled longitudinally (Riet), making her the only detectable recipient of new strains, yet no other Chimpanzee A members were sampled far enough in advance of Riet to identify as potential donors (Figure **1b**). Based on the sampling criteria for transmission events, our null distribution included 197 donor-recipient dyads for which we had the potential to detect transmission events.

Compared to the null distribution, microbial transmission events were strongly enriched between animals living in the same exhibit (binomial regression, coefficient= 1.8932, p<0.001). This may result from microbial transmission through conspecific-directed social behaviors, similar diets or gut physiology in animals of the same species, and/or limits on microbial dispersal distances (Figure **3b**). To examine each of these hypotheses in turn, we separately considered the sets of within- and across-exhibit transmission events. If social behaviors are the primary drivers of microbial spread in this system, we might expect the highest transmission rates within chimpanzees or within bonobos, which frequently engage in contact-based behaviors such as grooming and non-conceptive sexual behavior (Rodrigues and Boeving 2019; Brooker et al. 2025), and perhaps next highest within gorillas, which form cohesive groups and forage together in the wild, but are thought to engage less often in contact-based affiliative behaviors (Stokes 2004; Cooksey et al. 2025). We expected orangutans, which are semi-solitary and do not form stable social groups in the wild, to affiliate least with their group-mates in captivity and exchange the fewest microbial strains (Galdikas 1985). However, there were no significant differences in the observed rates of within-exhibit transmission (binomial regression; p=0.441) (Figure **3c**).

We next considered across-exhibit transmission events. Transmission across exhibits was not significantly enriched or depleted among phylogenetically close host species (binomial regression; coefficient=x, p=0.149) (Figure **3d**). Transmission across exhibits was significantly *lower*, not higher, between physically close exhibits as evaluated by the pairwise distances between the exhibit centers based on GPS data (binominal regression; coefficient=862.40, p=0.037). However, it is not clear whether the combined distribution of the indoor and outdoor exhibits accurately represents the animals’ locations during the sampling months (February-May), as they were mostly confined to the indoor portions during this time period. The indoor exhibits that were directly adjacent in the primate house did not have significantly more transmission (binomial regression, coefficient=-0.378, p=0.315).

### Comparisons to wild primates and humans

We considered that microbial transmission across exhibits might be mediated by humans: Zookeepers handling the food and supplies of multiple exhibits, for instance, or guests closely approaching multiple exhibits consecutively. If this were true, microbial species transmitted across exhibits might be more likely to be typically shared with humans, and less likely to be typically shared with wild great apes, compared to species transmitted within exhibits. To test this hypothesis, we drew on publicly available metagenomic data from a nearby human population (*Homo sapiens* from Germany) and from wild-living great apes, including wild gorillas (*Gorilla gorilla gorilla* from Republic of the Congo, Central African Republic, Gabon); wild bonobos (*Pan paniscus* from Democratic Republic of the Congo); wild chimpanzees (*Pan troglodytes schweinfurthii* from Uganda; *Pan troglodytes verus* from Côte d’Ivoire). No gut metagenomic samples were available from wild orangutans at this time (Figure **4a**).

**Figure 4:**
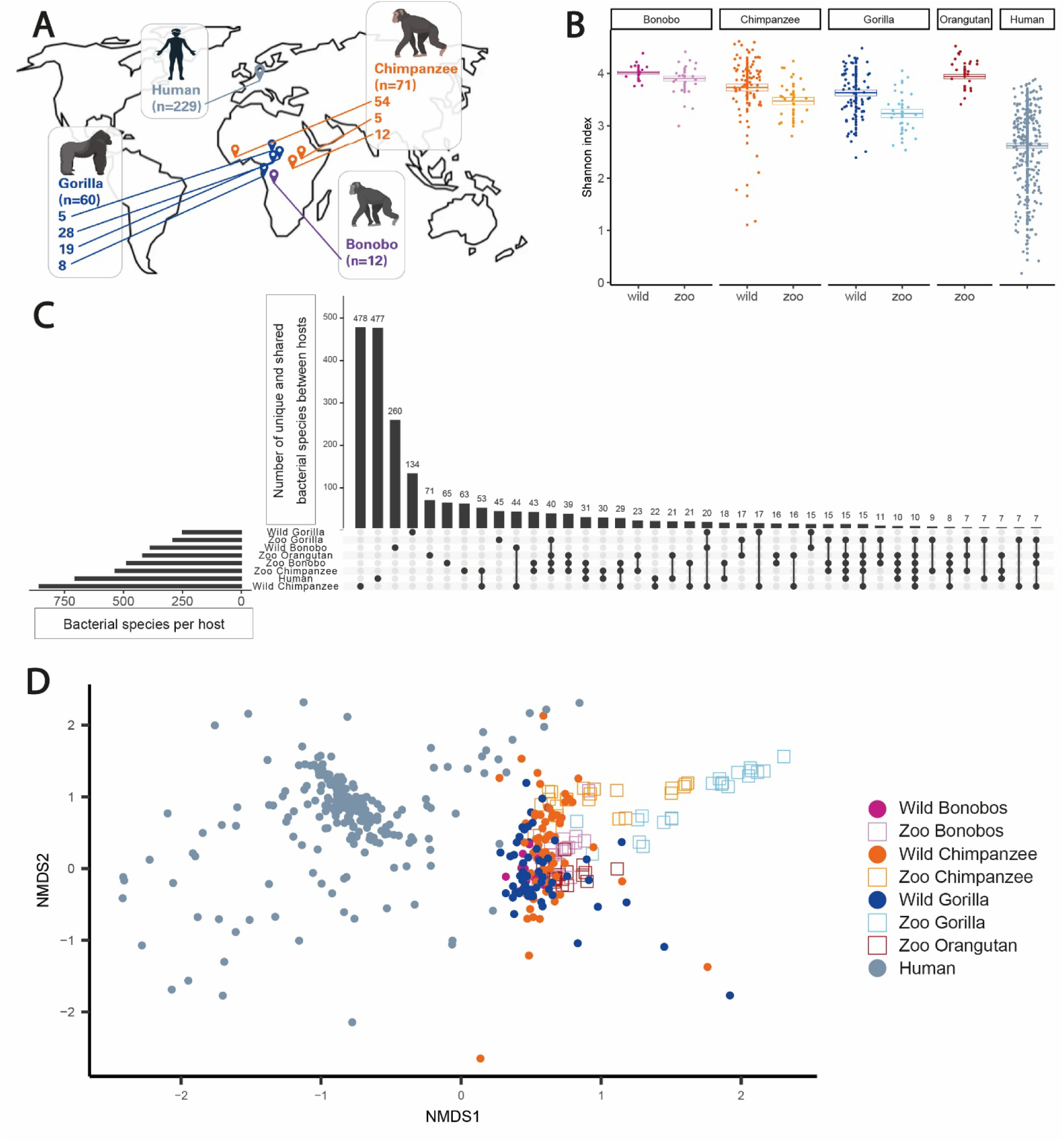
Microbiome of captive non-human great apes converge but are not humanized. **a**) Public gut metagenomic datasets from humans living in Tübingen, Germany and from wild great ape populations in several African locations. Whenever possible, multiple wild populations were included to represent the extent of natural microbiome variation. No publicly available metagenomes for wild orangutans were found. **b)** Shannon diversity of zoo animals, their wild counterparts, and humans. Gorillas differed the most between the wild and captivity, followed by chimpanzees and then bonobos (gorillas: F= 13.766, p= 0.0004; bonobos: F= 3.276, p= 0.080; chimpanzees: F= 3.950, p= 0.0499). **c)** Upset plot showing the datasets and the number of unique/shared bacterial species. Vertical bars show numbers of bacterial species, horizontal bars show sample size of each dataset and connected dots indicate sharing among datasets. **d**) NMDS ordination based on Bray-Curtis dissimilarities. Closer points indicate more similar microbiome composition while more distant points indicate less similar microbiome composition.

For chimpanzees, bonobos, and gorillas, we could directly compare microbiome composition between wild and captive conspecifics. Shannon diversity was generally lower in captive apes than in wild apes, though the magnitude of the difference varied by species (gorillas: F= 13.766, p= 0.0004; bonobos: F= 3.276, p= 0.080; chimpanzees: F= 3.950, p= 0.0499). Both wild and captive apes had more diverse gut microbiomes than humans (Figure **4b**). The majority of microbial species detected in the study were both host-species-specific and found only in the wild (478 species found in wild chimpanzees but no other populations; 260 found only in wild bonobos, 134 found only in wild gorillas). A smaller number of microbial species were shared across host species in the same setting (zoo or wild). For example, 40 microbial species were found in all four zoo species but in no wild populations, while 20 microbial species were found in all three wild species but never in the zoo. Others were host-species-specific regardless of the setting: for instance, 53 species were shared between zoo and wild chimpanzees, but were not present in any other wild or captive population. Humans tended to share species mainly with chimpanzees, regardless of their location (30 shared species with zoo-living chimpanzee and 22 with wild-living chimpanzee) but also with other zoo-living apes (with 15 shared species among all zoo-living animals, 21 with zoo-living orangutans, 18 with zoo-living bonobos, and 7 with zoo-living gorillas) (Figure **4c**).

In an NMDS ordination, captive chimpanzees and bonobos had similar microbiome compositions to wild chimpanzees and bonobos. Gorillas had the most distinct microbiomes from their wild counterparts, consistent with previous reports that the diets and gut microbiomes of folivorous primates are the most altered in captivity (Frankel et al. 2019). However, none of the great ape species in our study showed evidence of a shift from their wild counterparts towards humans in captivity (Figure **4d**).

### Microbial traits predicting transmissibility

We classified microbial species as either transmitted primarily across exhibits, transmitted equally across and within exhibits, or transmitted primarily within exhibits. To be reliably classified into one of these categories, the species had to be transmitted more than once. Then, the species was assigned to one of the categories depending on whether it was transmitted more often within or across exhibits or equally. Contrary to our initial expectation, species transmitted across exhibits were no more likely to be shared with humans than species transmitted within exhibits, as almost no species involved in transmission in the great apes were ever detected in humans. In fact, many transmitted species were typical of wild great ape gut microbiomes (Figure **5a**).

**Figure 5:**
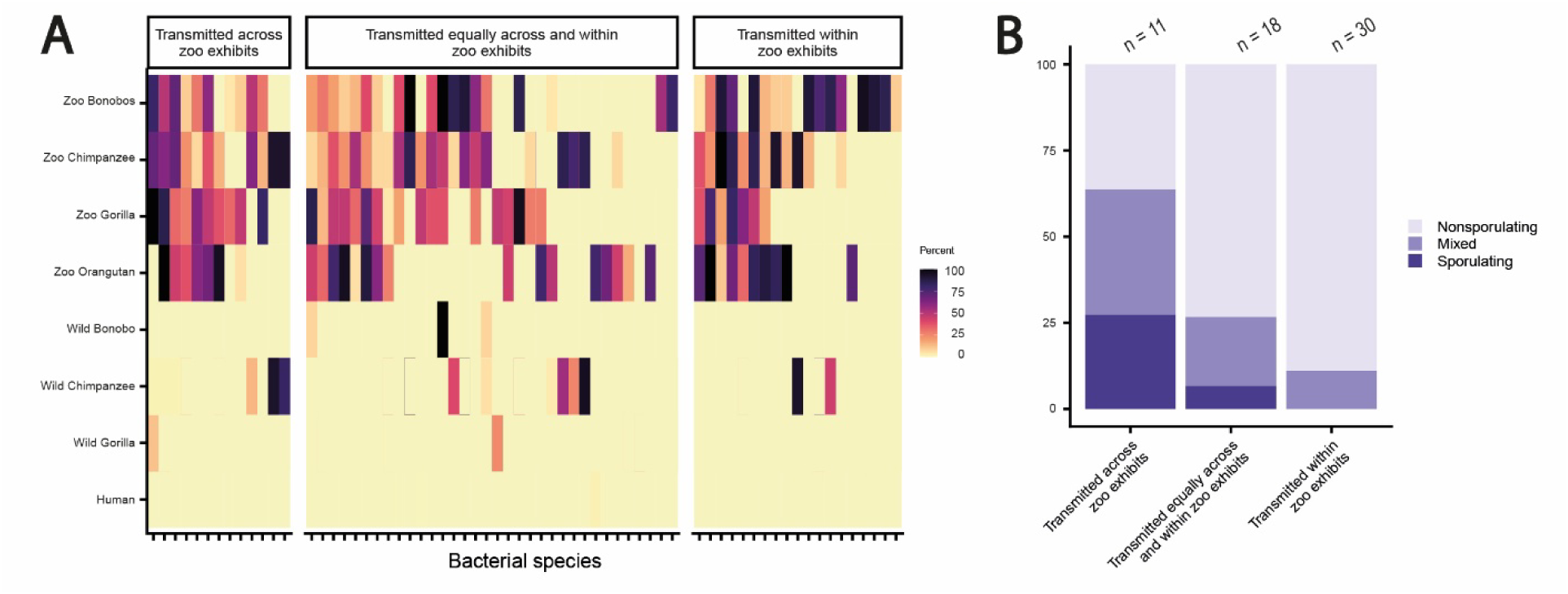
Bacterial dispersal ability, not presence in human gut microbiomes, predicts transmission between different zoo exhibits. Bacterial species involved in multiple transmission events in the zoo were categorized as transmitted across zoo exhibits, transmitted equally across and within zoo exhibits, or transmitted within zoo exhibits. **a**) Heat map represents all bacterial strains that fall into one transmission category. Each tile represents the percentage of samples per dataset the microbial species was found in. Darker colors indicate higher percentages. **b**) Bacterial species assigned to transmission category were further classified as belonging to spore-forming genera, non-spore-forming genera, or genera with a mix of sporulating and non-sporulating species. Species transmitted mainly across exhibits were more likely to come from spore-forming genera.

We next annotated aerotolerance and spore formation properties on a genus-level basis for each species involved in transmission, as both have been linked to increased probabilities of long-distance transmission (Valles-Colomer et al. 2023). Aerotolerance did not vary with transmission mode, as the vast majority of gut microbial species were predicted to be obligately anaerobic. However, microbial species transmitted across exhibits were significantly more likely to come from spore-forming genera than species transmitted within exhibits (cumulative linked mixed model, coefficient= −2.760, p= 0.0047) (Figure **5b**).

## DISCUSSION

Many studies have reported patterns of phylosymbiosis and host-microbe co-diversification among wild great ape populations (Sanders et al. 2023; Narat et al. 2020; Nishida and Ochman 2021; Rühlemann et al. 2024). However, it is often difficult to tell whether co-diversification patterns are driven by adaptation of bacterial strains to certain host species or geographic constraints on bacterial transmission (Good 2026). Zoo-based primate research offers the opportunity to monitor multiple species in a common environment, while still having species-typical social groups and pseudo-natural conditions. Zoos often bring together different animal species that would likely not encounter each other in wild settings, allowing us to see whether microbial populations that are geographically restricted in the wild are capable of colonizing other host species when given the opportunity. We combined broad sampling of many individuals to compare strain sharing within and between exhibits, with opportunistic longitudinal sampling of available individuals to investigate patterns of new strain acquisitions.

Even in a shared environment with controlled diets, strong host-species-specific patterns emerged in microbiome composition that mirrored the host species phylogeny. Because the Zoo Leipzig has two separately housed chimpanzee groups, we could partially disentangle the separate contributions of group living and host species identity. Strain sharing levels between chimpanzees in different exhibits fell between those of animals of the same species living together and animals of different species living apart, indicating that spatial constraints on transmission and then differential survival of bacteria in different host species both contributed to microbiome variation across exhibits. Within exhibits, the most similar samples were typically those collected from the same individual over time. While individual microbiome uniqueness is a common observation in wild animal and human studies (Chen et al. 2021; Björk et al. 2022), it may be surprising to find such strong individual signatures in the zoo as well, where animals are confined to a smaller area with all their group-mates and have presumably had many opportunities to exchange microbes over the years. Possible host attributes that could maintain individuality in shared enclosures could include age, wild or captive birth, dominance hierarchies that influence food access within groups, and exposure to antibiotics (Torfs et al. 2025).

We tracked new strain acquisitions that occurred during the study period and identified their potential donors. New strains were much more likely to come from another member of the same exhibit than a member of a different exhibit, similar to previous work showing that group membership constrains the transmission of bacterial strains in wild baboons (Debray et al. 2026). There were no substantial differences in the rates of transmission within exhibits despite the variation in social behaviors among great ape species (Rodrigues and Boeving 2019; Brooker et al. 2025; Stokes 2004; Cooksey et al. 2025; Galdikas 1985), potentially because the zoo housing imposes artificial constraints on social behaviors, such as bringing together orangutans into closer proximity than they would normally live in the wild. Our strain tracking analysis suggested that bacterial transmission was possible across exhibits and occurred independently with respect to host species phylogeny, indicating that phylogenetic structure only emerges in the longer term as strains establish and survive with different success in different species. Dispersal across exhibits could occur through the air or with the assistance of animal vectors such as birds, insects, or humans (zookeepers) that interact with multiple habitats in succession. The enrichment of spore-forming bacteria in cross-exhibit transmission suggests airborne transmission of spores as a likely pathway.

Finally, we compiled and analyzed publicly available metagenomic sequences from wild gorillas, bonobos, and chimpanzees, as well as humans living in Germany (approximately 500 km from the Zoo Leipzig). Each of the captive apes clustered near their wild counterparts in an NMDS ordination, indicating that host-species-specific microbiome differences are at least partially maintained in captivity. However, the majority of microbial species that were restricted to only one ape species in the wild were lost from their captive counterparts. This observation was consistent with previous work (Nishida and Ochman 2021) and suggests that these bacteria may be particularly specialized, e.g., on particular components of great ape diets in the wild that are not available in captivity. Unlike some previous studies (Clayton et al. 2016; Houtz et al. 2021), we found no evidence that zoo-housed apes have more human-like microbiomes than their wild counterparts. Species sharing with humans constituted a minor fraction of the great apes’ microbiomes overall, and microbial species transmitted between zoo exhibits were almost never detected in humans. Interestingly, a previous study of saliva microbiomes of the Zoo Leipzig chimpanzees and bonobos also noted that captive apes clustered with their wild counterparts rather than with humans (J. Li et al. 2013), suggesting that exposure to humans and their microbiomes may be a zoo-specific phenomenon. Certain practices such as diet and enclosure design might influence the extent of microbiome humanization. The great apes living at the Zoo Leipzig eat diets high in vegetables and raw leaves, with very little processed food. Enclosures are large and in many cases, physical barriers such as rivers keep visitors at a distance from the animals.

One limitation of this study is that sampling occurred only between February and May, showing a snapshot of one season. The microbiomes of wild great apes often vary seasonally depending on food availability (Hicks et al. 2018) and we also might expect small changes as the diet plans of the Zoo Leipzig varies slightly by seasons. During the sampling time animals were mainly housed inside, which might influence transmission opportunities within and across exhibits. We additionally note that because fecal samples were opportunistically collected, exhibits varied in their sampling coverage and some individuals were not sampled at all. This limits our ability to definitively pinpoint the donors of new strain acquisitions. We can only identify one or more possible donors, acknowledging that the true donor may have been unsampled, and that we may miss some strain transmission events that newly appeared in recipients but were not traceable to any of the sampled animals. Lastly, we sampled only the animals’ feces, but sampling the objects and surfaces within and between their habitats could help to resolve transmission intermediates. Our annotation of bacterial characteristics suggests survival on surfaces as a key driver of long-distance transmission, but future work that explicitly samples hosts and their environments will be highly valuable as well.

In this study, both spatial constraints on bacterial transmission and host-species-specific bacterial adaptation appear to shape the gut microbiome composition of a community of captive living great apes. Transmission was shaped by group housing (i.e., within versus between exhibits) and bacterial ability to form spores. Whether new strains are retained, however, appears to depend on host filtering, as phylogenetic structure emerged in strain- and species-level composition in the long term. Finally, we found that bacteria can disperse among nearby exhibits of great ape species that would never encounter one another (or presumably one another’s microbiomes) in the wild. The implications of this finding for primate health are not known, and future research should further address the welfare consequences of inter-species microbial transmission in captivity.

## METHODS

### Animal information and sample collection

All study subjects were housed in indoor-outdoor enclosures in the Pongoland center at the Zoo Leipzig (see **Table S1** for full list of subjects and group membership). During the months of sample collection (February to May), they were primarily confined to their indoor enclosures due to the cooler temperatures. Animals were fed an omnivorous diet composed of leaves (lettuce, cabbage, chicory, and other foliage), fruits (pear, kiwi, orange, banana, grape, apple, melon), vegetables (leek, celery, broccoli, cauliflower, fennel, cucumber, red pepper, onion, turnip, carrot, tomato, beetroot), boiled meat, boiled egg, and yogurt. Lettuce and vegetables were provided on a daily basis in the mornings, fruit was provided on a daily basis in the evenings, and meat, eggs, and yogurt were provided weekly. Older and low-ranking individuals (who have low priority of access in their social groups) were hand-fed in the mornings and evenings to ensure that they received adequate nutrition. Older or underweight individuals sometimes get additional cooked potatoes or noodles.

Fecal samples were collected from sleeping cages or during behavioral testing sessions led by the Wolfgang Köhler Primate Research Center. Feces were stored in 95% ethanol in a 5°C cooling room for up to five days, then transported to the Max Planck Institute for Evolutionary Anthropology, where they were frozen at −20°C.

### Metagenomic sequencing

We extracted DNA from fecal samples using the Qiagen PowerSoil kit, then prepared metagenomic libraries using the SeqWell PurePlex DNA Library Preparation Kit. Libraries were sequenced on a NovaSeq X 10B flow cell to a median depth of 20 million reads/sample. Metagenomic shotgun reads were trimmed to remove adapter sequences and for quality using *trimmomatic* v0.39, with the parameters: *trimmomatic PE -threads 16 ILLUMINACLIP:NexteraPE-PE.fa:2:30:10 SLIDINGWINDOW:4:20 MINLEN:70*. After, host reads were removed using *bowtie2* v2.5.4 (Langmead and Salzberg 2012) with the parameter -p 20, by aligning the reads against the respective host genome: human - GRCh37, bonobo - panPan3, gorilla - gorGor6, chimpanzee - panTro6, orangutan - ponAbe3. Endogenous (host) DNA content in fecal samples ranged from 0.39% to 6.5% of the total metagenomic reads.

### Species- and strain-level profiling

To determine species-level composition of each sample *kraken2* v 2.1.6 was used for taxonomic classification (Wood et al. 2019), followed by *bracken* v3.0.1 to estimate relative abundance of microbial species (Lu et al. 2017). For strain-level composition *inStrain* v1.9.0 and *SynTracker* v1.4.0 were used in parallel, as both apply different approaches for strain-level evaluation. When running *inStrain* we used default parameters (99.999% ANI and 25% of the microbial genome covered in both samples at 5x or greater coverage required to call a shared strain). Alongside *SynTracker* was employed after the reads were assembled into contigs using *megahit* v1.2.9 (D. Li et al. 2015). The contigs were passed to SynTracker, where we required an average pairwise synteny score (APSS) of 95%. The proportion of reads that mapped to microbial genomes in our database ranged from 28-65 % per sample, varying slightly by host species. Chimpanzee samples had the highest alignment rates (32-65%), followed by bonobos (46-64%), orangutans (44-57%), and gorillas (28-45%).

### Spatial layout of exhibits

We collected spatial information by traversing the entire outer boundary of Pongoland, all inner walkways, and the indoor primate house, recording GPS points every 7-10 steps. We summarized spatial distances between exhibits by calculating the following metrics: i) A binary variable (yes/no) describing whether each pair of exhibits shared any open boundaries, either because their outdoor enclosures were directly adjacent or because the open sections of their indoor enclosures were directly across the hallway from each other; ii) The pairwise distance between the centroids of each indoor/outdoor exhibit area.

### Comparisons with wild populations and humans

To compare zoo-living animals to their wild counterparts and humans, we analyzed publicly available metagenomic datasets. The public datasets encompass: gorillas (*Gorilla gorilla gorilla* from Republic of the Congo (Campbell et al. 2020; Hicks et al. 2018), Central African Republic (Amato et al. 2019), Gabon (Rühlemann et al. 2024)); bonobos (*Pan paniscus* from Democratic Republic of the Congo (Rühlemann et al. 2024)); chimpanzees (*Pan troglodytes schweinfurthii* from Uganda (Amato et al. 2019; Rühlemann et al. 2024); *Pan troglodytes verus* from Côte d’Ivoire (Rühlemann et al. 2024)); and humans (*Homo sapiens* from Germany (Suzuki et al. 2022)). See **Table S2** for accession numbers of microbiome datasets.

Wild samples were analyzed using the same pipelines as the zoo samples (see *Metagenomic sequencing* and *Species- and strain-level profiling*), with the same parameters. Briefly, quality control started with *trimmomatic* and host removal using *bowtie2*. After, species-level composition was assessed using *kraken2* and *bracken*, and *inStrain profile*. Alpha and beta diversity metrics were calculated based on bacterial species with at least 500 reads per sample, and after rarefying samples to 100,000 reads.

### Annotating microbial phenotypes

We searched the Joint Genome Initiative’s Genomes OnLine Database for lists of known anaerobic, aerobic, and facultative bacteria. For each reference genome in our microbial database, we extracted the GOLD information on all bacterial species from the same genus. In most cases, a genus was composed of all obligately anaerobic species (n=1673 cases), all obligately aerobic species (n=46 cases), or all facultatively aerobic species (n=30 cases). The remaining genera were coded with mixed annotations, e.g., “aerobic/anaerobic”. Similarly, we extracted lists of spore-forming and non-spore-forming bacteria and coded each reference genome as sporulating, non-sporulating, or mixed according to the phenotypes of all other bacterial species from the same genus.

### Controls, replication, and statistical analysis

We originally generated technical replicates at the DNA extraction stage (i.e., independent DNA extraction from the same fecal sample) from 24 of the 58 samples to assess repeatability of the sequencing workflow. At the outset of the analyses in this paper, we selected one replicate per biological sample at random to carry forward to all downstream analyses to prevent pseudoreplication and ensure that significance tests would accurately reflect the power of the study. Each set of 18 samples was sequenced alongside a commercial DNA positive control (ZymoBIOMICS Microbial Community DNA Standard II) and a DNA extraction blank.

Variation in strain sharing was modeled using linear regression with either the dyad type coded as a categorical variable (same individual, different individuals in the same exhibit, etc.) or estimated divergence time between host species coded as a continuous variable. Transmission rates were modeled by constructing a null distribution of all possible donor-recipient pairs for which enough samples were available to detect transmission if it had occurred, then running a binomial regression with the outcome variable representing whether or not we had actually observed transmission between each donor-recipient pair. Shannon diversity in captive gorillas, bonobos, and chimpanzees was compared to each host species’ wild counterpart using species-specific ANOVA models (note that no metagenomic sequences from wild orangutans were available at this time). We assessed the relationship between transmission mode and bacterial aerotolerance using a cumulative linked mixed model, which are designed to handle ordinal data. Each bacterial species could fall into one of three ordered categories: from a genus entirely composed of non-oxygen-tolerant species, from a genus composed of a mix of oxygen-tolerant and non-oxygen-tolerant species, or from a genus composed entirely of oxygen-tolerant species. Similarly, we tested the relationship between transmission mode and bacterial spore formation based on whether each bacterial species came from a genus entirely composed of sporulating species, from a genus composed of a mix of sporulating and non-sporulating species, or from a genus composed entirely of sporulating species. Both aerotolerance and spore formation displayed relatively strong phylogenetic structure, with a minority of genera falling into the “Mixed” category.

### Research ethics

All fecal sample collection was opportunistic, non-invasive, and did not interfere with the daily routine of the study animals. The procedure was approved by the Zoo Leipzig and complies with the European Association of Zoos and Aquaria (EAZA) Minimum Standards for the Accommodation and Care of Animals in Zoos and Aquaria, the World Association of Zoos and Aquaria (WAZA) Code of Ethics and Animal Welfare, and the Association for the Study of Animal Behavior and Animal Behavior Society (ASAB/ABS) Guidelines for the Treatment of Animals in Behavioral Research and Teaching. Institutional Animal Care and Use Committee (IACUC) review was not necessary to conduct this research because it did not meet the criteria for animal experiments under the German Animal Welfare Act.

## Notes

### Competing Interest Statement

The authors have declared no competing interest.

